# Comprehensive comparison of large-scale tissue expression datasets

**DOI:** 10.1101/010975

**Authors:** Alberto Santos, Kalliopi Tsafou, Christian Stolte, Sune Pletscher-Frankild, Seán I. O’Donoghue, Lars Juhl Jensen

**Author notes:** Corresponding authors: Alberto Santos, Addr: Blegdamsvej 3b, DK-2200 Copenhagen N, Lars, Juhl Jensen, Addr: Blegdamsvej 3b, DK-2200 Copenhagen N. These authors contributed equally (Alberto Santos, Kalliopi Tsafou). Current affiliation: Ferring Pharmaceuticals, Copenhagen, Denmark (Sune Pletscher-Frankild). Email addresses: AS, KT, CS, SPF, SIO, LJJ.

## Abstract

For tissues to carry out their functions, they rely on the right proteins to be present. Several high-throughput technologies have been used to map out which proteins are expressed in which tissues; however, the data have not previously been systematically compared and integrated. We present a comprehensive evaluation of tissue expression data from a variety of experimental techniques and show that these agree surprisingly well with each other and with results from literature curation and text mining. We further found that most datasets support the assumed but not demonstrated distinction between tissue-specific and ubiquitous expression. By developing comparable confidence scores for all types of evidence, we show that it is possible to improve both quality and coverage by combining the datasets. To facilitate use and visualization of our work, we have developed the TISSUES resource (http://tissues.jensenlab.org), which makes all the scored and integrated data available through a single user-friendly web interface.

## Introduction

Mapping out which proteins are present in each tissue is of major importance for understanding the functional differences between tissues as well as their development and differentiation (Pontén et al., 2009; Emig & Albrecht, 2011). Several high-throughput experimental technologies have been used for this, the most widely used of which are expressed sequence tags (ESTs) (Wheeler, 2003; “UniGene: A Unified View of the Transcriptome,” 2003), high-density oligonucleotide microarrays (also called DNA chips) (Su et al., 2004; Clark et al., 2007), and RNA sequencing (RNA-seq) (Krupp et al., 2012; Fagerberg et al., 2014; Uhlen et al., 2015).

ESTs are short sequence reads — typically around 400bp — derived from 5’ or 3’ ends of complementary DNA (cDNA) libraries from tissues or cell lines (Adams et al., 1991; Bailey, Searls & Overton, 1998; Nagaraj, Gasser & Ranganathan, 2007). Consequently, for a highly expressed gene, one would expect to see a correspondingly high abundance of ESTs derived from its transcripts. A more recent sequencing-based approach to quantifying transcript levels is RNA-seq. The major difference to EST sequencing is that random cDNA fragments are sequenced instead of only the 5’ and 3’ ends. The resulting reads are aligned to a reference genome, producing a quantitative expression profile for each gene (Wang, Gerstein & Snyder, 2009; Nagalakshmi, Waern & Snyder, 2010). Because reads are generated from all parts of a transcript instead of only the ends, the number of reads observed for a gene depends on both its length and its level of expression. A major advantage of RNA-seq is the ability to resolve individual splice variants if enough reads are obtained for a gene. Microarrays are another extensively used technology for transcriptome analysis. Gene expression is quantified by measuring the fluorescence intensity of labeled cDNA that hybridizes to oligonucleotide probes (Lipshutz et al., 1999; Harrington, Rosenow & Retief, 2000; Churchill, 2002). Because a microarray can contain millions of different probes, the transcript levels of all genes can be measured simultaneously.

The above mentioned techniques are all based on measuring mRNA levels. Fewer techniques exist for high-throughput measurement of protein levels. One of them is multiplexed immunohistochemical staining of tissue samples embedded in paraffin blocks (sometimes referred to as tissue microarrays). Histological analysis of the resulting images of tissues stained with an antibody can semiquantitatively tell where the target protein is present (Kampf et al., 2012). The main challenge to using this approach at the proteome scale is the need for specific antibodies against all proteins (Buchwalow et al., 2011). Mass spectrometry has also been used for measuring protein abundances in tissue samples, mainly in bodily fluids (Adkins, 2002; Schmidt & Aebersold, 2006; Aretz et al., 2013), muscle biopsies (Lundby et al., 2012), and tumor samples (Schwartz, 2004; Seeley & Caprioli, 2008; Paul et al., 2013). Two recent publications collected many of these experiments into a single repository (Wilhelm, Schlegl & Hahne, 2014) and for the first time used this technology for in-depth proteomic profiling of a broad selection of normal human tissues (Kim et al., 2014).

Large-scale tissue expression datasets have formed the basis for many analyses and discoveries related to correlations between data from different technologies, mainly between transcriptomics and proteomics experiments (Waters, Pounds & Thrall, 2006; Bitton et al., 2008), roles of housekeeping and tissue-specific genes in protein complexes (Bossi & Lehner, 2009; Emig & Albrecht, 2011), biological processes (Zhu et al., 2008b; Chang et al., 2011; Schaefer et al., 2013), and diseases (Shyamsundar et al., 2005; Vasmatzis et al., 2007; Lage et al., 2008; Magger et al., 2012; Börnigen et al., 2013). However, the majority of these studies (Shyamsundar et al., 2005; Lage et al., 2008; Bossi & Lehner, 2009; Chang et al., 2011; Magger et al., 2012; Schaefer et al., 2013; Börnigen et al., 2013) are based solely on microarray data from the GNF Expression Atlas (Su et al., 2004), which could bias the results. It is thus relevant to test to which extent the different technologies and datasets give congruent results.

We here present the first comparative evaluation of the quality of tissue associations from a variety of different datasets and experimental methods as well as from manual curation (The UniProt Consortium, 2014) and automatic text mining of the biomedical literature (Figure 1). We show that these datasets — despite the technological differences — agree surprisingly well with each other and can be combined to improve quality and coverage. Finally, as a result of the integration process, we have developed the TISSUES resource (http://tissues.jensenlab.org), which makes the above mentioned heterogeneous data more easily accessible to researchers by collecting them in a single place and assigning confidence scores.

**Figure 1.**
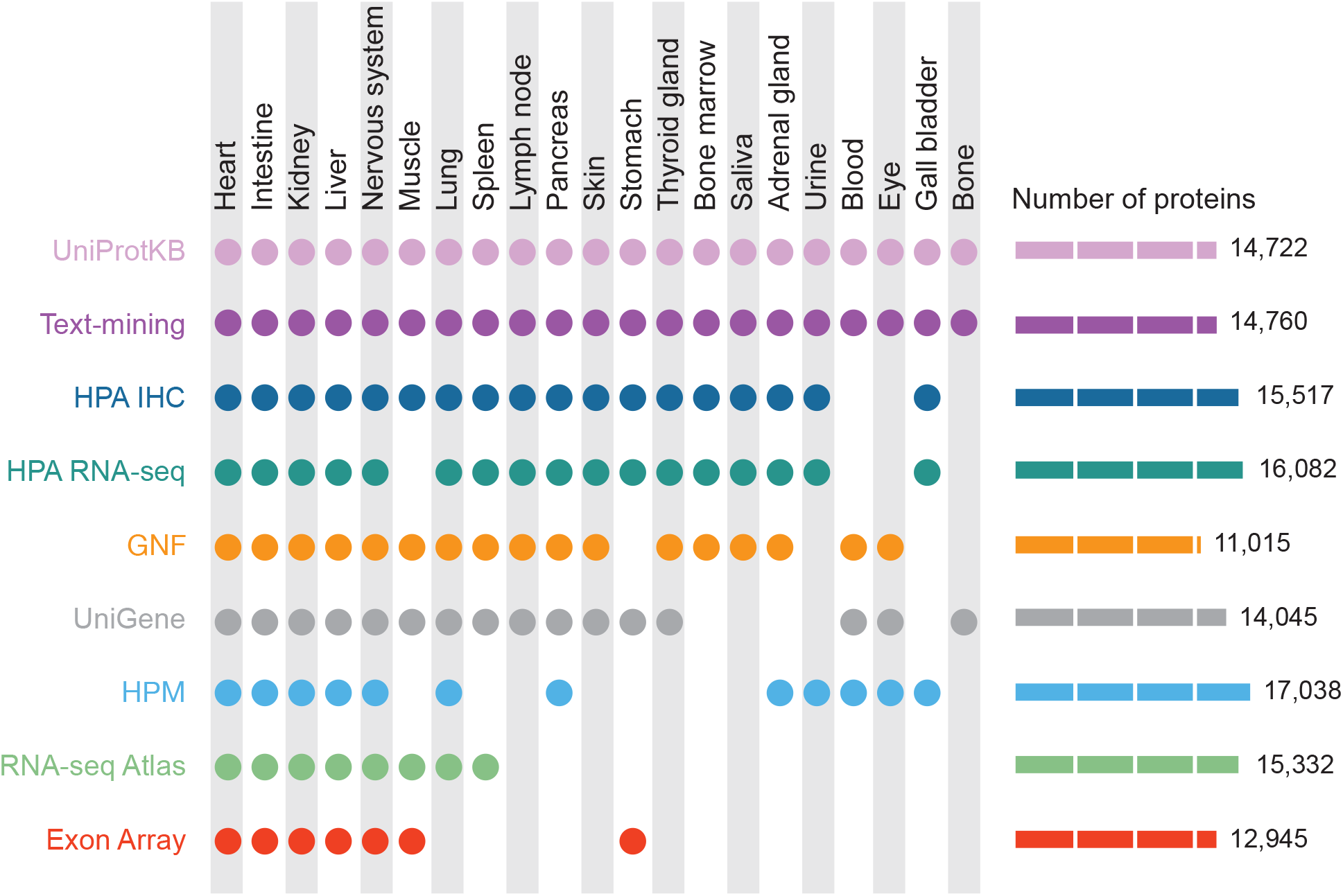
Summary of the tissues and number of proteins present in each dataset. For our analyses, we mapped 9 datasets to 21 major tissues of interest. This figure shows which datasets cover which of these major tissues and how many proteins each dataset identified.

## Methods

### Datasets

#### GNF Gene Expression Atlas

The experimental data from the Human U133A/GNF1H Gene Atlas) (Su et al., 2004) was downloaded from the BioGPS portal (http://biogps.org/). The dataset contains information for 44,775 probe sets, which we filtered to remove probe sets associated with multiple targets (names ending with “_[r,i,f,x]_at” and control probe sets (names starting with “AFFX”). We mapped the remaining probe sets to gene identifiers using the probeset-to-gene annotation file (gnf1h.annot2007.tsv) and finally mapped these to 16,598 Ensembl protein identifiers using the alias file from the STRING database (Franceschini & Szklarczyk, 2013). The GNF Gene Expression Atlas provides information for 79 tissues, 60 of which we could map to Brenda Tissue Ontology terms. We scored each gene–tissue association based on the normalized expression units obtained from the microarray analysis, under the assumption that transcripts identified with higher intensity are less likely to be false positives. When multiple probe sets mapped to the same gene, we used the mean expression value.

#### Affymetrix Exon tiling array

These high-density microarrays (Clark et al., 2007) contain probe sets for more than one million annotated and predicted exons. We downloaded the data from the Gene Expression Omnibus (Barrett et al., 2011) (GSE5791 series matrix) and used the 565,690 probe sets mapped to a gene identifier according to the GPL4253 platform. We mapped the latter to 15,559 Ensembl protein identifiers. The Exon Array experiment examined 16 tissues mainly from the nervous system studying six sub-regions of the brain. All tissues could be mapped to BTO terms. As in the other microarray experiment, we used the mean normalized expression units as the score for each gene–tissue association.

#### UniGene

The UniGene database (Wheeler, 2003; “UniGene: A Unified View of the Transcriptome,” 2003) clusters together Expressed Sequence Tags (EST) that belong to a single gene and includes information about the tissue where each EST was observed. We used the *Homo sapiens* UniGene Build #236, which contains 24,289 clusters that could be mapped to 18,493 Ensembl protein identifiers via the provided gene symbols or UniGene cluster identifiers. UniGene Human library (Hs.lib.info) provides information for 80 tissues from which we discarded several with ambiguous names, e.g. “retina and testis” or “uncharacterized tissue” (Data S1), and finally obtained 60 BTO terms. The scoring scheme for UniGene is based on the number of ESTs clustered into a single gene that belong to the same tissue. When multiple clusters mapped to the same gene, we used the total number of ESTs from the clusters.

#### RNA-seq atlas

The RNA-seq Atlas (Krupp et al., 2012) is a web-based resource that provides expression data for 21,399 genes in 11 tissues. We mapped the genes to 18,063 Ensembl protein identifiers using the STRING alias file; all the specified tissues mapped directly to BTO terms. We used the normalized Reads Per Kilobase per Million mapped reads (RPKM) as the confidence score for each gene–tissue association.

#### HPA RNA-seq data

The Human Protein Atlas version 12 (Fagerberg et al., 2014; Uhlen et al., 2015) provides short-read high-throughput sequencing data (RNA-seq) in 27 non-disease tissues. We mapped 20,315 Ensembl gene identifiers for which the database contained expression levels to 18,491 Ensembl protein identifiers and all the tissue names to BTO terms. Similarly to the scoring scheme applied to the RNA-seq Atlas dataset, we assigned the normalized expression levels in Fragments Per Kilobase of exon per Million fragments mapped (FPKM) as the confidence score for each gene–tissue association.

#### HPA Immunohistochemistry

HPA also provides an atlas of protein expression derived from immunohistochemistry experiments over many tissues (Fagerberg et al., 2014; Uhlen et al., 2015). We obtained information on the expression of 16,384 genes in 45 tissues (data downloaded on 21st January 2014), which we mapped to 15,552 Ensembl Protein identifiers and 45 BTO terms. For each antibody and tissue, HPA provides a semiquantitative strength of staining (*staining*_*a,t*_), which we translated into numeric values (not detected: 0, low: 1, medium: 3, high: 6). When only a single antibody was used to measure a protein, we simply used the staining values from that antibody as the confidence scores for the tissues.

When multiple antibodies for the same protein were used we used a more complex scoring scheme to combine the staining values from the individual antibodies:

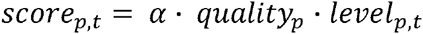

where α is a scaling factor for making the multi-antibody scores comparable to the single antibody scores, *quality*_*p*_captures the internal agreement among the antibodies for the protein, and *level*_*p,t*_ is a weighted average of staining values of the antibodies for the protein in a given tissue.

The correction factor for the quality of the antibodies is defined as:

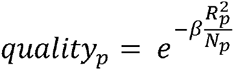

where *β* is a parameter optimized as described below, *N*_*p*_ is the number of antibodies for the protein, and *R*^2^ measures the disagreement between the antibodies across all tissues:

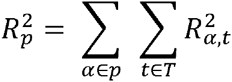

where T is the set of tissues studied and 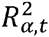 is the disagreement in a given tissue between one antibody and the average of the antibodies:

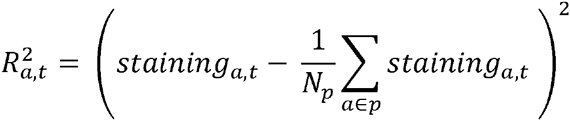

We defined the level of a protein in a given tissue (*level*_*p,t*_)as a weighted average of the antibodies:

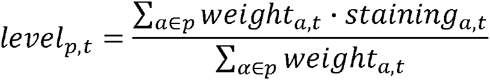

where the weights are defined based on the disagreements between the antibodies:

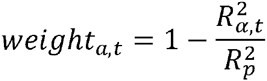

We validated the scoring scheme and determined the values of the free parameters by calculating the fold enrichment (see Quality of proteomics data) against UniProtKB. The optimal values of the parameters were *α* = 30 and *β* = 0.7.

#### Human Proteome Map

HPM is a large mass spectrometry-based catalogue of protein profiles in 30 normal human tissues (Kim et al., 2014), which contains more than 290,000 tryptic peptides. We mapped these to Ensembl by comparing the sequences to all theoretical tryptic peptides derived from Ensembl v75 protein sequences, allowing for up to two missed cleavages. We assigned each tryptic peptide to the corresponding Ensembl gene identifier and mapped these to a total of 17,038 Ensembl protein identifiers using the STRING alias file. The 30 normal human tissues were comprised of 17 adult tissues, 7 fetal tissues, and 6 primary hematopoietic cell types. Because the corresponding adult and fetal tissues map to the same term in BTO, the 30 tissues mapped to only 26 different BTO terms. For those tissues mapping to the same BTO term, we averaged the number of tryptic peptides. As confidence score for a protein being expressed in a given tissue, we used the unique number of tryptic peptides observed.

#### UniProtKB tissue annotations

UniProtKB (The UniProt Consortium, 2014) provides manually curated protein annotations. This includes annotations of tissue expression for 17,075 human proteins. Whereas each protein is typically only annotated with one or a few tissues, the number of different tissue terms used is very high; we were able to manually map UniProtKB tissues for 401 different BTO terms in total. Because the annotations are manually curated, we considered all protein–tissue associations from UniProtKB to be of the highest confidence.

#### Text mining

The text mining pipeline used in this work has been described in detail elsewhere. It relies on an efficient dictionary-based named entity recognition algorithm (Pafilis et al., 2013) and a co-occurrence scoring scheme (Binder et al., 2014) to extract associations from Medline abstracts. To use the pipeline to extract of protein–tissue associations, we complemented the existing dictionary of human gene and protein names from STRING with a dictionary of tissue and cell types constructed from BTO. The pipeline extracted more than one million protein–tissue associations based on co-occurrences of 16,748 proteins and 5,300 BTO terms.

### Evaluation and calibration of scores

To evaluate the quality of the gene–tissue associations from each dataset, we compared them to the UniProtKB gold standard. We quantified the agreement in terms of the fold enrichment, which we define as the fraction of pairs in a dataset that are also in the gold standard divided by the fraction expected by random chance. The latter is defined as the fraction of possible gene–tissue pairs that are found in the gold standard. For these fold-enrichment calculations we considered only the genes and tissues that are shared between the dataset and the gold standard.

We calculated the fold enrichment for score windows of 100 gene–tissue associations to capture the relationship between fold enrichment and the quality scores defined in the previous sections. To be able to convert the quality scores from individual datasets into confidence scores that are comparable between datasets, we first fit the relationships between quality scores and fold enrichments with mathematical functions with only a few parameters. We used these to define the low-, medium-, and high-confidence cutoffs for the comparisons of the datasets (Table 1). Next, we performed a global transformation of the fold enrichments into the “star” confidence scores used in the COMPARTMENTS resource (Binder et al., 2014) based on the text-mining scores, which the two resources have in common. The combined, calibrated functions for translating quality scores into the final confidence scores are listed in Table 1 (Figure S3).

**Table 1.**
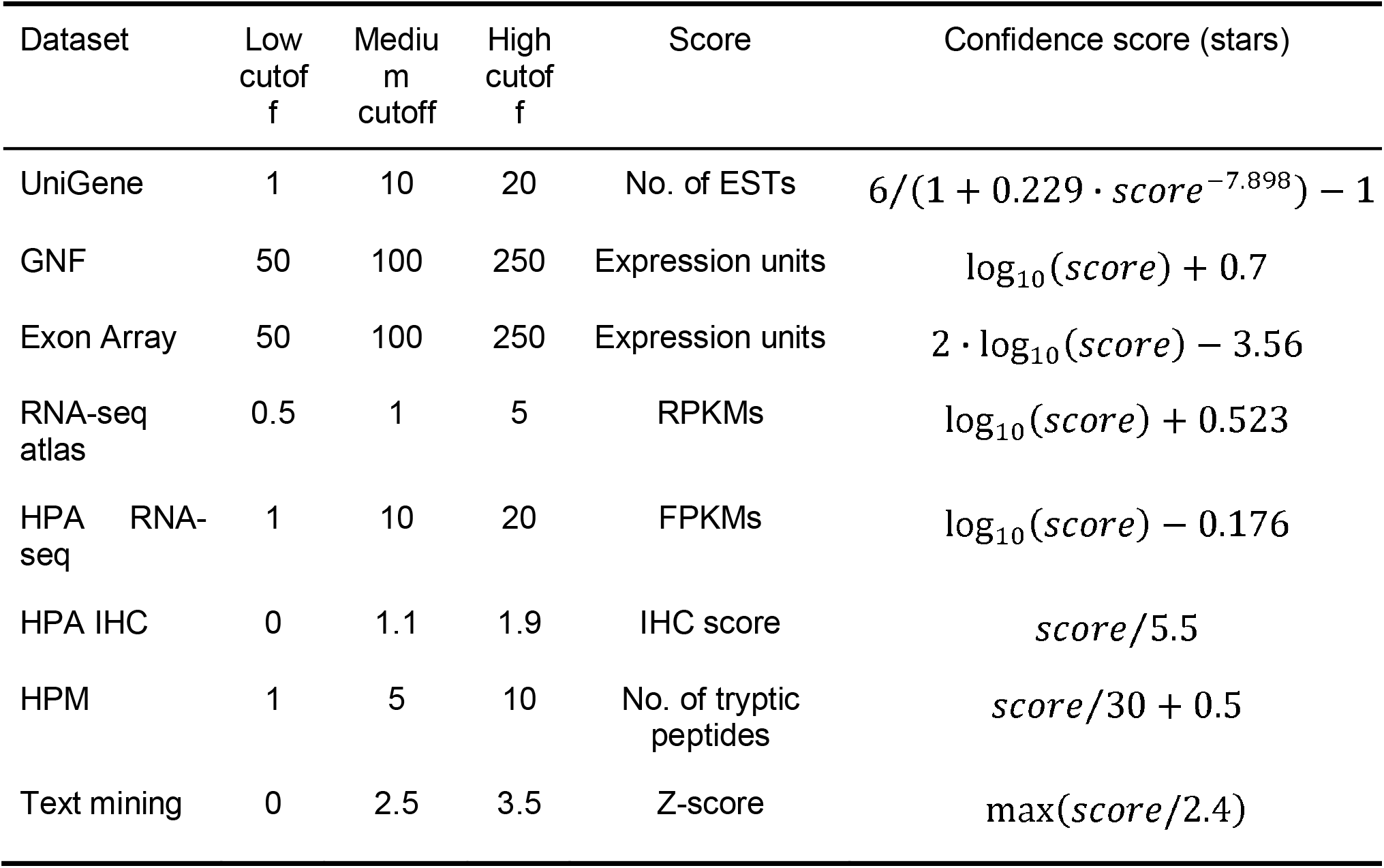
*Definition of cutoffs.* This table shows the different confidence cutoffs used in the analyses for each dataset, the quality score and how each quality score is converted to the unified confidence score used in the TISSUES web resource.

### Web resource

To make the protein–tissue associations available for query by a web resource, we store all data in a PostGreSQL database. The web interface is implemented through the same Python web framework used for the COMPARTMENTS database (Binder et al., 2014). The body map onto which the data is visualized was manually created in Adobe Illustrator and saved as a Scalable Vector Graphics (SVG). In the user’s browser, JavaScript is then used to provide interactive coloring and labeling of tissues.

## Results and Discussion

To systematically compare the different datasets, we standardized the varying names used for the same tissues to their respective terms in the Brenda Tissue Ontology (Schomburg et al., 2013) (Data S1). Because this ontology is structured as a directed acyclic graph, this also helps deal with the challenge of different datasets having different tissue resolution; for example, some datasets study the brain as a whole whereas others study different parts separately. We decided to base our analyses on the 21 major tissues shown in Figure 1.

### Tissue-specific and ubiquitous transcripts

Many studies have made the distinction between housekeeping genes, which are expressed in most tissues, and tissue-specific ones, which are expressed in only a few tissues (Hsiao et al., 2001; Lercher, Urrutia & Hurst, 2002; Liang et al., 2006; Dezso et al., 2008; Zhu et al., 2008a,b; Eisenberg & Levanon, 2013). However, there are no strict definitions of these two classes of genes, and it is not clear to what extent this represents a natural classification. To answer the latter, we analyzed the expression breadth of five transcriptome datasets, i.e. how many genes are expressed in how many tissues. As this depends strongly on the threshold used to decide whether a gene is expressed in a given tissue, we performed the analysis with three different cutoffs, in the following referred to as low, medium, and high confidence (see Methods).

Figure 2 shows the expression breadths for five transcriptome datasets, each at the three different confidence levels. Most show a clear bimodal distribution with peaks at the extreme ends, i.e. the vast majority of genes are expressed either in only a few tissues or in most tissues measured. We thus show that data from several sources and technologies robustly support a natural distinction between tissue-specific and ubiquitously expressed genes.

**Figure 2.**
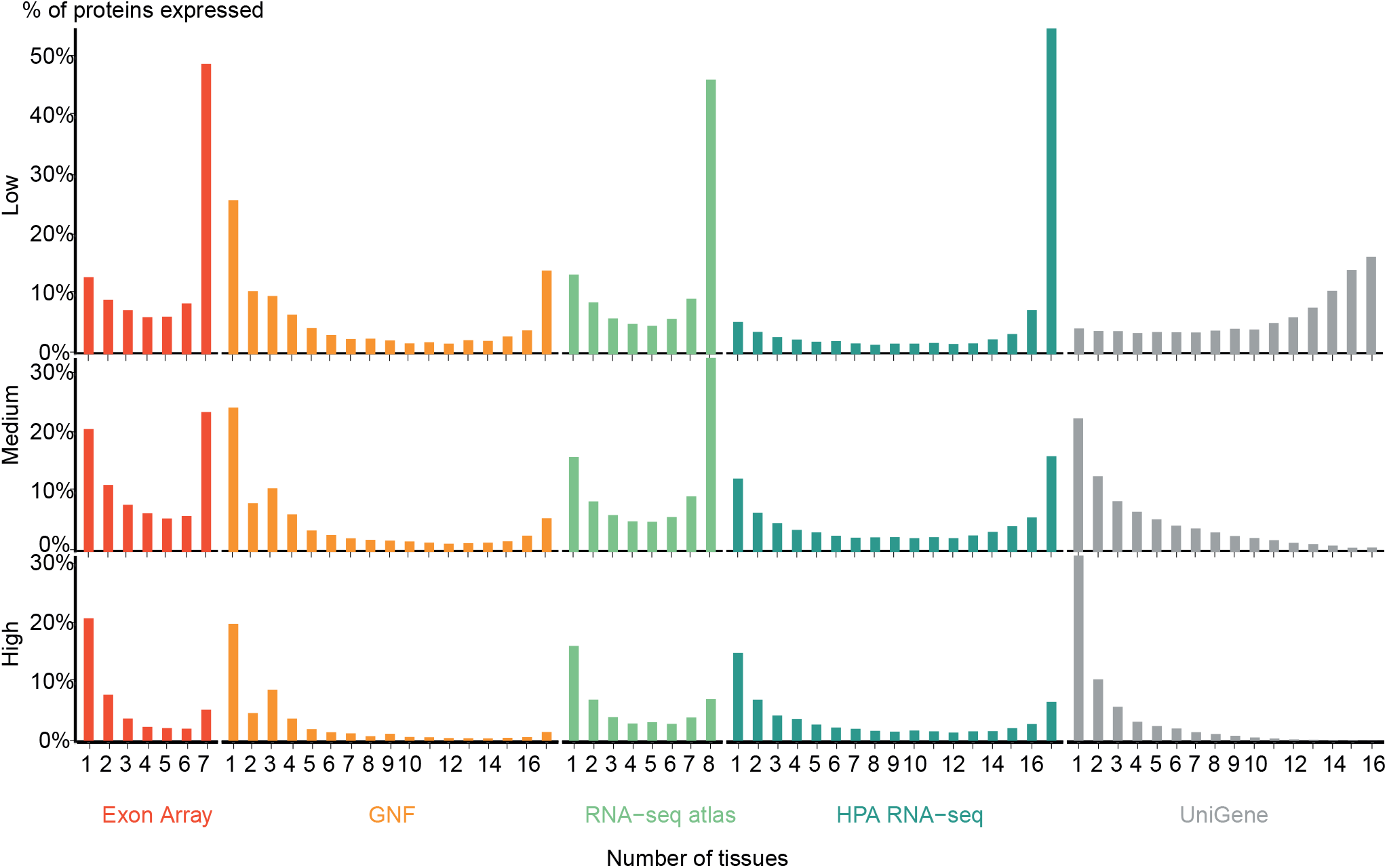
Distribution of expression breadth of the transcriptome datasets. For each of the five mRNA datasets, the histograms show the number of protein-coding genes expressed at low, medium, and high confidence as function of number of tissues. With the exception of UniGene, the distributions are bimodal, with most proteins occurring in either few tissues or in most tissues measured, supporting the notion of distinguishing between tissue-specific and ubiquitous expression.

Zhu and colleagues (Zhu et al., 2008a) also showed a bimodal trend when comparing the GNF expression atlas and EST sequencing data; however, for the latter data type the bimodality was weak. We similarly find very few tissues-specific genes when analyzing UniGene at the low-confidence cutoff, but show that this trend is reversed when using more stringent cutoffs. We observe that the GNF dataset is atypical in that it identifies fewer ubiquitously expressed genes at all cutoffs than the rest of the datasets, including the other microarray-based study (Exon array).

### Consistency of transcriptomic methods

The previous analysis showed that the global trends in terms of tissue specificity are similar across the transcriptome datasets. That, however, does not imply that the datasets necessarily agree on which genes are expressed where. To quantify the agreement, we focused on the five tissues and 3,254 genes covered by all the transcriptome datasets. Comparing the five transcriptome datasets, we saw that genes are assigned to tissues with high consistency between datasets at all three confidence levels (Figure 3) (P < 10^-15^ for all pairwise overlaps). At medium confidence 39.2% (5679/14504) of gene–tissue associations are common to all datasets and 65.8% (9537/14504) are common to at least four of the five datasets (Data S2).

**Figure 3.**
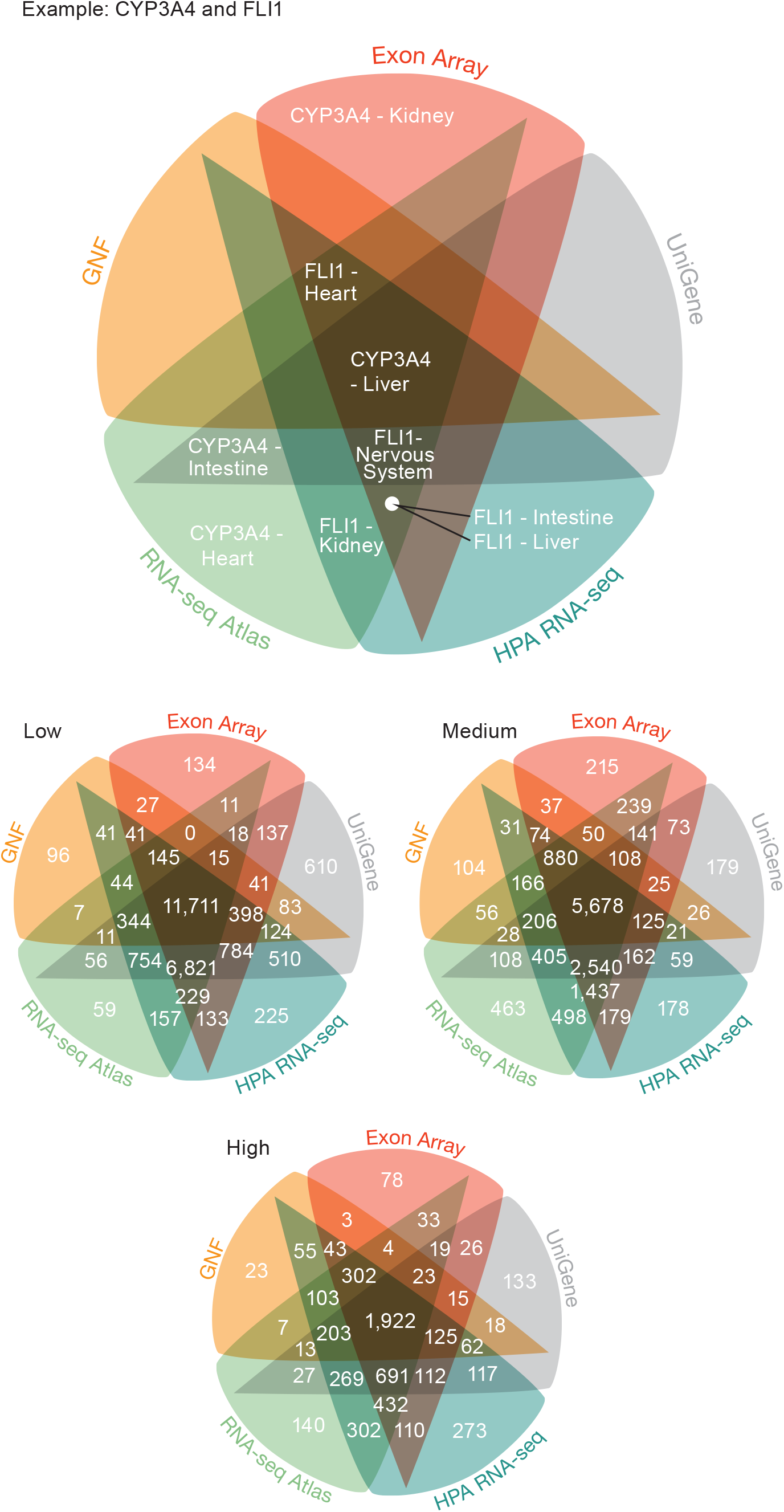
Consistency of the transcriptome datasets. We assessed the consistency of the five transcriptome datasets by calculating the overlap of gene–tissue associations for the shared genes and tissues. At all levels of confidence, we observe surprisingly good agreement, with the largest count in each Venn diagram representing associations found by all five datasets.

The largest discrepancy in the comparison is the large number of gene–tissue associations found by all datasets except GNF at all three confidence levels (Figure 3). This is likely because the GNF Expression Atlas was made using microarrays designed prior to the completion of the Human Genome Project, which consequently have suboptimal probe sets for many genes.

Conversely, the largest agreement is seen among the three most recent datasets, which were generated using RNA-seq or exon arrays. At medium confidence, their overlap makes up 72.65% (10538/14504) of all gene–tissue associations, 13.66% (1439/10538) of which are not found by any other dataset.

### Correlation between expression values and confidence levels

The high consistency between the mRNA datasets demonstrates their quality; however, it does not guarantee that the selected cutoffs are comparable and represent the same level of confidence across datasets. To assess the assumed correlation between expression values and confidence, we compared all datasets to a gold standard of gene–tissue associations extracted from scientific literature by UniProtKB (The UniProt Consortium, 2014). While reliable, UniProtKB annotations are very incomplete as they are restricted to what has been published. It is thus not possible to estimate the precision of a dataset; instead, we quantified the quality of the datasets in terms of its fold enrichment of correct gene–tissue associations compared to random chance.

The comparison showed that fold enrichment for gold-standard associations increased steadily with expression value from all datasets (Figure 4A). This was expected because, in general, the more abundant a transcript, the more reliably it can be identified. Moreover, we find that the low-, medium-, and high-confidence cutoffs used in the preceding analyses correspond to the same quality in all datasets. However, a dataset of lower quality will give fewer associations at any given confidence cutoff.

**Figure 4.**
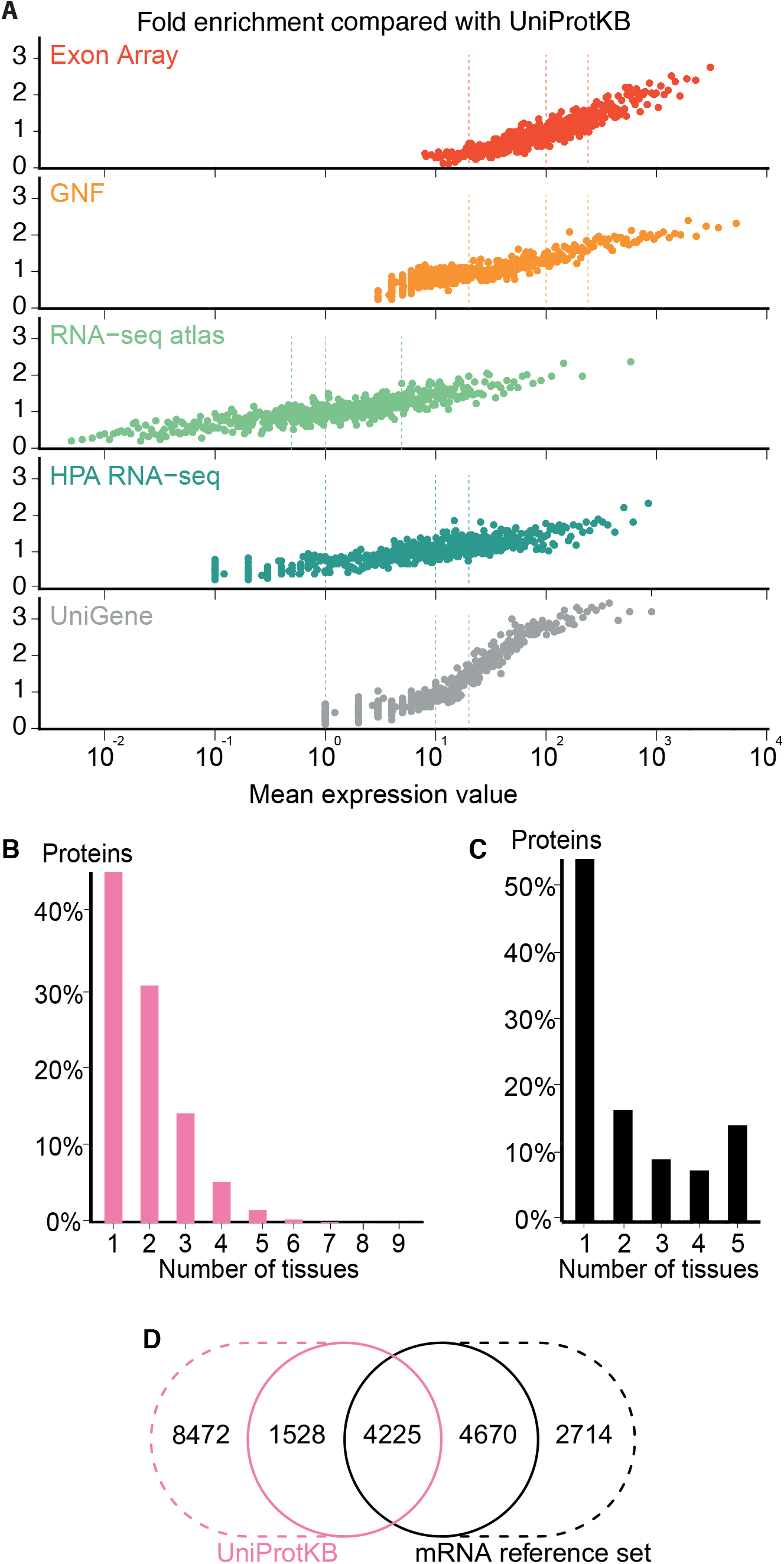
Quality of the transcriptome datasets. *A.* To assess the correlation between expression level and confidence, we compared the transcriptome datasets to a gold standard, namely UniProtKB. We quantify the quality of the datasets in terms of its fold enrichment for correct gene–tissue associations compared to random chance. The comparison shows that higher expression values imply higher quality and that the three confidence cutoffs (vertical dotted lines) used correspond to equivalent quality in all datasets. *B.* The distribution of expression breadth for UniProtKB is strongly skewed towards tissue-specific proteins, contrary to what was seen for transcriptome datasets. We thus constructed a consensus mRNA reference set; its expression breadth distribution is in line with that of the individual mRNA datasets. *D.* The mRNA reference set is highly complementary to the UniProtKB gold standard, providing 7,384 gene– tissue association that are not in the latter.

The expression breadth distribution of UniProtKB is strongly skewed towards tissue-specific proteins; only 0.72% of proteins (106/14722) are annotated as expressed in more than five tissues. This reflects that many annotations describe proteins as widely or ubiquitously expressed but list only a few tissues. Also, UniProtKB annotations are incomplete, because many proteins have only been described in the literature as present in some of the tissues where they are expressed.

In light of this and the high quality of the mRNA datasets, we built a complementary set of gene–tissue associations, hereafter called the mRNA reference set, with high-confidence support from at least three datasets. This set exhibits the expected bimodal distribution of expression breadth (Figure 4C) and provides 7,384 gene–tissue associations not present in UniProtKB (Figure 4D, Data S3).

### Quality of proteomics data

To complement the mRNA datasets with protein-level data, we investigated the Human Protein Atlas immunohistochemistry data (HPA IHC) (Fagerberg et al., 2014; Uhlen et al., 2015) and the mass spectrometry data from the Human Proteome Map (HPM) (Kim et al., 2014). To compare these with other datasets, we developed a quality scoring scheme for each as described in the methods section.

With the scoring schemes defined, we analyzed the two proteomics datasets with respect to enrichment for associations from both the UniProtKB and mRNA reference sets (Figure 5A). Higher scores were correlated with higher enrichment, validating that the proposed scoring schemes work. Despite looking at proteins instead of transcripts, the proteomics datasets show worse fold enrichment than the transcriptome datasets, when compared to the UniProtKB gold standard. This is consistent with the criticism raised over the quality of the HPM data based on an analysis of olfactory receptors expressed in multiple tissues (Ezkurdia & Vázquez, 2014), which demonstrated a high percentage of false positives in this dataset. In case of HPA IHC, this is especially true for data derived based only on a single antibody.

**Figure 5.**
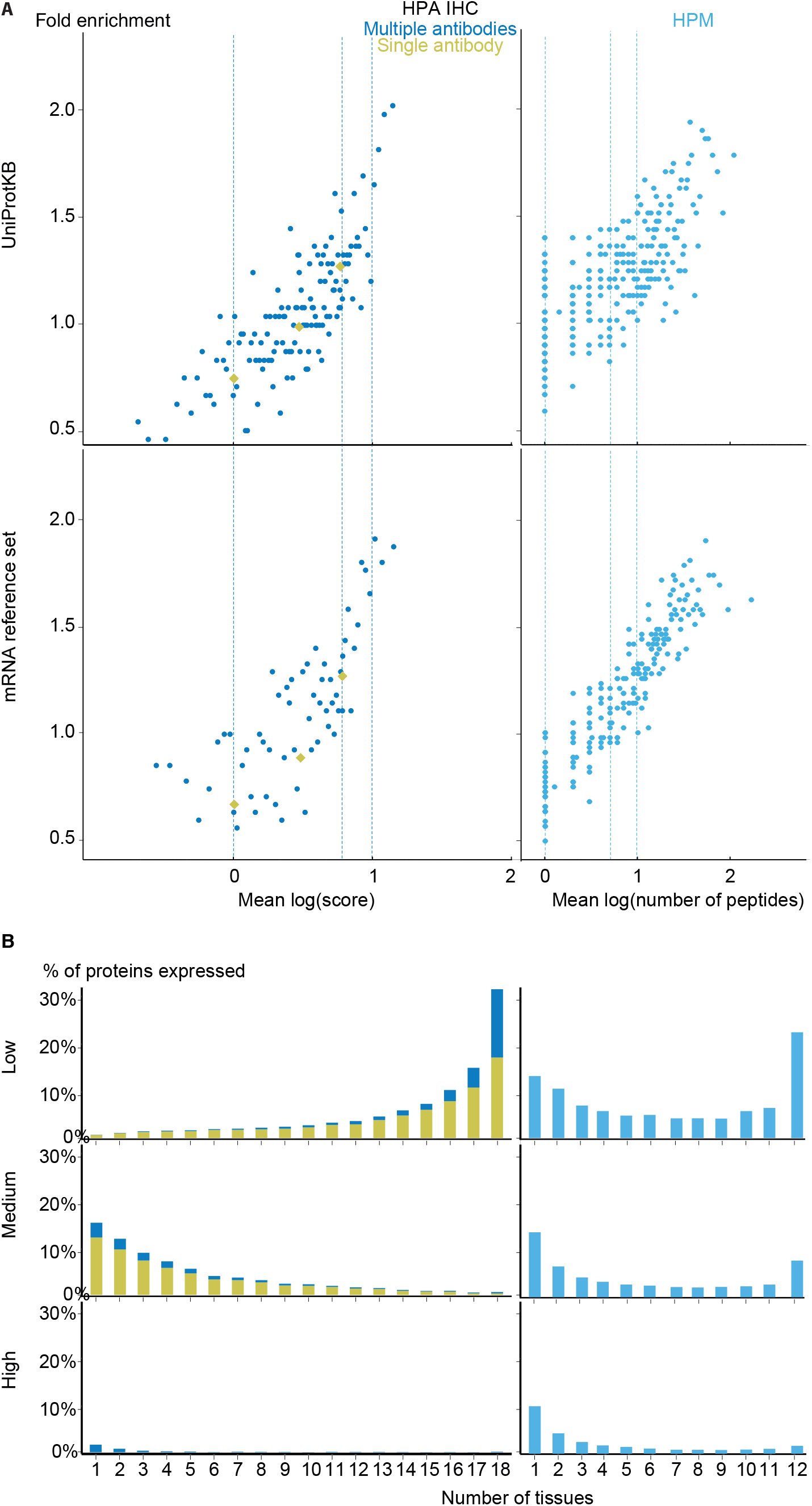
Analysis of the proteomic datasets. *A.* To make the data from HPA IHC and HPM comparable with other datasets, we developed a quality scoring scheme for each. The quality scores show good correlation with the fold enrichment for associations from the UniProtKB and the mRNA reference sets. *B.* The distribution of expression breadth is consistent with the results of the transcriptome datasets in case of HPM, whereas the results for HPA IHC vary qualitatively between confidence levels.

HPM exhibits bimodal distributions of expression breadth at all confidence levels (mainly at low and medium levels) consistent with the majority of the transcriptome datasets (Figure 5B). This consistency across confidence levels is in part due to a substantial fraction (18,027/106,021) of the associations from HPM being high confidence. Conversely, the HPA IHC dataset is dominated by low-confidence associations for proteins studied with only a single antibody or with multiple antibodies that gave different results. At low confidence, proteins tend to be associated with many tissues, which is likely due to unspecific antibodies. By contrast, most proteins have higher-confidence links to only a few tissues.

### Complementary annotations from text mining

Automatic text mining of the biomedical literature has the potential to extract information that has been either overlooked by curators, not yet curated, or not annotated due to curation standards (Aerts et al., 2008; Van Auken et al., 2012). We used a previously published text-mining pipeline (Pafilis et al., 2013; Binder et al., 2014), expanded with a dictionary of tissues and cell lines, to extract associations between genes/proteins and tissues and scored them according to their co-occurrence in sentences and abstracts.

We evaluated the quality of these associations by comparing them to both the UniProtKB and mRNA reference sets (Figure S1A). This analysis shows that co-occurrence-based text mining performs well for this task. The high agreement with UniProtKB is not surprising considering that text mining and curation are both based on the available literature. The comparison to the mRNA reference set, however, shows that many of the associations found by text mining, but not by curators, are also supported by direct experimental evidence.

The distribution of expression breadths is, like for UniProtKB, skewed towards the tissue-specific end (Figure S1B), due to the same literature limitations. However, text mining associates each gene/protein with more tissues, even at high confidence. For example, 421 are linked to more than five tissues, which is four times more than what UniProtKB annotates. These results demonstrate the value of complementing manual annotation with automatic text mining.

### Improved tissue profiles through data integration

So far we have shown that the quality of the different datasets is comparable at each of the chosen confidence levels. To assess the consistency and complementarity of different data sources, we compared the medium-confidence associations from UniProtKB and text mining to two pooled sets of high-confidence associations from transcriptomics and proteomics experiments, respectively.

Despite the inherent differences between data types and technologies compared, when looking at the common proteins and tissues, 43.4% (17,053/39,294) of all associations are supported by at least two of the four sets (Figure 6A). The transcriptomics and proteomics sets show the largest pairwise agreement, which accounts for 32.12% (11,472/35,709) of the associations from the two sets and 29.2% (11,472/39,294) of all associations (Data S4). This agreement highlights the strong connection between transcription and final protein abundance; indeed, transcription was recently demonstrated to explain about 80% of the differences seen in protein expression (Li, Bickel & Biggin, 2014).

**Figure 6.**
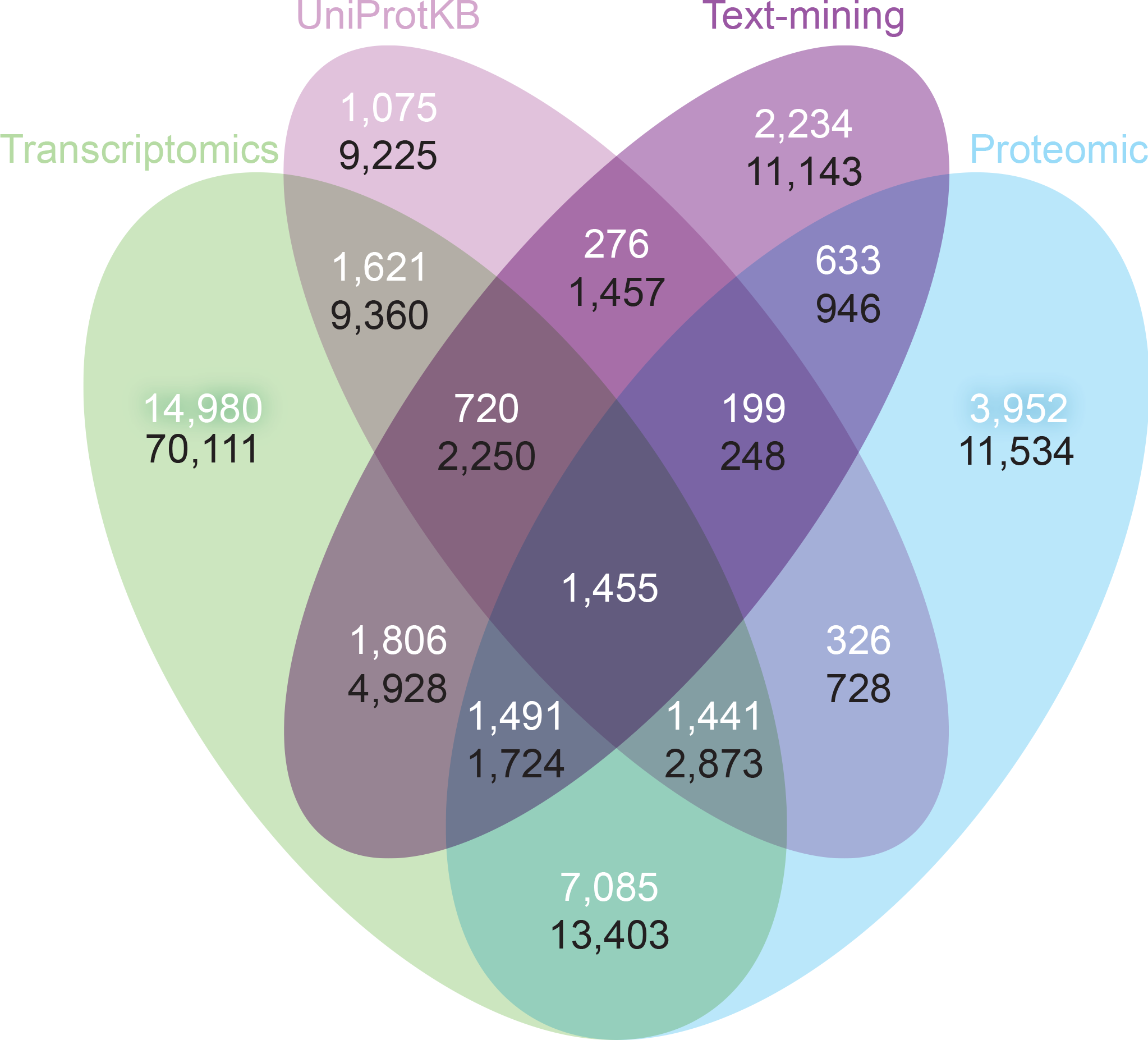
Consistency and complementarity of evidence types. To assess the consistency and complementarity of the associations supported by different types of evidence, we compared the medium-confidence associations from UniProtKB and text mining to two pooled sets of high-confidence associations from transcriptomics and proteomics experiments, respectively. The white numbers show the overlap of protein– tissue associations when considering only at the common proteins and tissues among all sets. The black numbers show the overlap when not restricting the comparison to common proteins and tissues. Together these analyses show that the different sources of evidence have high consistency across the common proteins and tissues, but that they are at the same time complementary because they cover different proteins and tissues.

Although all the sets are consistent on the proteins and tissues they have in common, they are also highly complementary because they cover different proteins and tissues. When not restricting the comparison to common proteins and tissues, 72.1% (102,013/141,385) of all the reported associations are unique to a single set (Figure 6B, Data S4). The analysis also reveals that only 6.5% (9,225/141,385) of the associations are unique to UniProtKB. Text mining alone captures 19.6% (5,410/27,596) of the curated literature results and complements them with 18,741 additional protein–tissue associations, 40.5% (7,598/18,741) of which are supported by the transcriptomics or proteomics sets.

Another way to illustrate the complementarity of the datasets is to compare the quality and coverage obtained when integrating many datasets compared to using a single dataset. To this end, we looked at the union of the transcriptomics and proteomics sets and compared it to the same number of top-scoring associations from the GNF atlas. Focusing on the 7,445 proteins and 17 tissues that GNF and UniProtKB have in common, 76% (11,395/14,974) of the associations from the integrated list were annotated in UniProtKB, whereas this was only the case for 60% (8,913/14,974) of the associations from GNF. Moreover, the integrated list includes 11,721 associations not covered by GNF (Data S5 and Figure S2).

### The TISSUES web resource

In light of the clear advantages of combining multiple datasets, we believe the scientific community can benefit from having a resource that integrates and provides easy access to the available information on tissue expression. We thus developed the TISSUES web resource that is available at http://tissues.jensenlab.org. Several other resources provide gene–tissue associations, including TiGER (Liu et al., 2008), BioGPS (Wu et al., 2009), TissueDistributionDB (Kogenaru et al., 2009), VeryGene (Yang et al., 2011), EBI Gene Expression Atlas (Kapushesky et al., 2010), and the GTEx portal (Lonsdale et al., 2013). What makes TISSUES unique is that it integrates data from many different technologies and sources, quantifies the reliability of each gene–tissue association, and thereby makes results from different sources comparable.

The web interface allows the user to search for a human gene and get a complete overview of where it may be expressed. To provide an at-a-glance overview, we show a body map with each the 21 major tissues colored according to the confidence that the gene of interest is expressed there (Figure 7). The figure also allows the user to see which sources of evidence support expression in a given tissue. Three interactive tables below the body map provide the user with more detailed information for the evidence from UniProtKB, high-throughput experiments, and text mining. This includes information on additional tissues, linkout to the source of the evidence whenever possible, and a unified confidence score ranging from 1 to 5 stars (see Methods).

**Figure 7.**
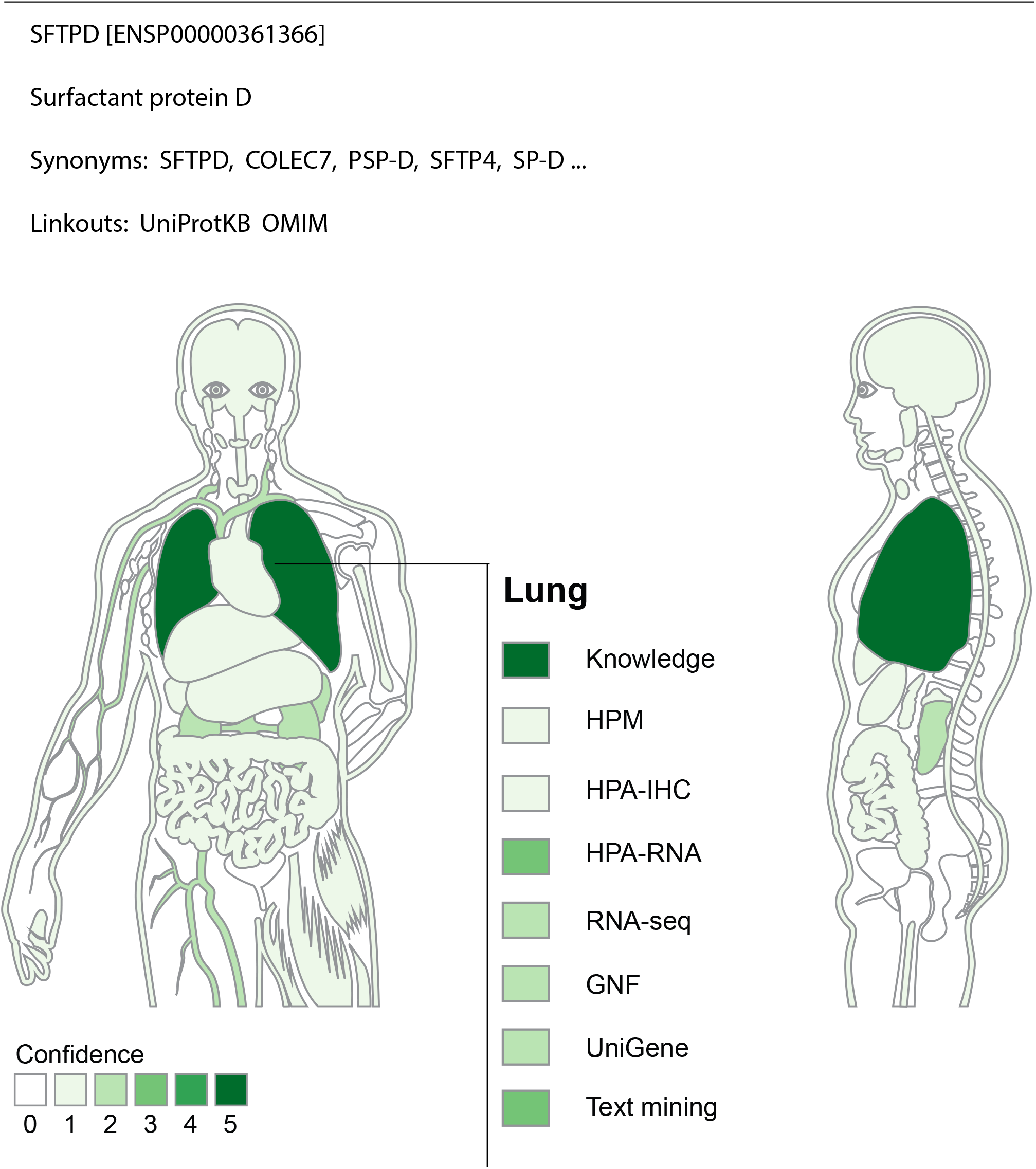
TISSUES: all data accessible in a single resource. The TISSUES web resource integrates all the data compared in this study, quantifies the reliability of each gene–tissue association, and thereby makes associations from different sources comparable. When searching for a human protein, the user is presented with a body map that provides a complete overview of where the protein is likely expressed by coloring the 21 major tissues according to the confidence of the protein–tissue association. The body map is interactive and allows the user to see which sources of evidence support expression in a given tissue. The TISSUES web resource is available at http://tissues.jensenlab.org.

TISSUES holds information for 21,294 genes and 5,305 different tissues and provides more than 2.2 million gene–tissue associations at varying confidence levels. These are all available for download under the Creative Commons Attribution License at http://tissues.jensenlab.org to facilitate large-scale studies.

## Acknowledgements

The authors thank Janos X. Binder for help with the web resource.

## Supporting Information

The data and the code used to get the fold enrichment analyses and to generate all the figures in the manuscript can be downloaded at:

https://github.com/albsantosdel/TISSUES-database_analyses

http://dx.doi.org/10.6084/m9.figshare.1409446

**Data S1 – Mapping of tissue names to Brenda Tissue Ontology terms.**

*This excel file contains the mapping from the tissue names from the original sources to the standardized BTO terms.*

*http://dx.doi.org/10.6084/m9.figshare.1405677*

***Data S2 - Common associations transcriptomic methods***

This file contains the following information:

- The list of genes studied at the different cutoffs
- The list of common associations to all datasets at the different cutoffs
- The list of common association for at least 4 datasets at the different cutoffs

*http://dx.doi.org/10.6084/m9.figshare.1405678*

***Data S3 - mRNA reference set associations***

This excel file contains the gene–tissue associations that form the mRNA reference set used in the fold-enrichment analysis.

*http://dx.doi.org/10.6084/m9.figshare.1405666*

***Data S4 - Common and unique gene–tissue associations to all the sets***

This file contains:

- Overlap between all the sets (transcriptomic set, UniProtKB, Text-mining and proteomics set)
- Overlap between the transcriptomic and the proteomic set
- The list of gene–tissue associations unique to each set

*http://dx.doi.org/10.6084/m9.figshare.1405679*

***Data S5 - Gene–Tissue associations coverage and quality analysis***

This file contains:

- Gene–tissue associations from the integration of the transcriptomics and proteomics datasets
- GNF atlas gene–tissue associations used in the analysis
- Overlap between the integrated set and UniProtKB
- Overlap between the GNF atlas studied set and UniProtKB

*http://dx.doi.org/10.6084/m9.figshare.1405680*

**Figure S1. Complementary annotations from text-mining** *A.* We used text-mining to extract associations between genes/proteins and tissues and score them based on their co-occurrence in sentences and abstracts. Comparing these associations to the UniProtKB and mRNA reference sets showed both the expected high agreement with UniProtKB and that many of the text-mined associations not annotated by curators are nonetheless supported by experimental evidence. *B.* The distribution of expression breadth for text mining is subject to the same literature limitations as UniProtKB. However, text mining associates each gene/protein with more tissues than the latter, even at high confidence, which demonstrates the value of complementing manual annotations with automatic text mining.

*http://dx.doi.org/10.6084/m9.figshare.1405672*

**Figure S2. Quality and coverage.** This figure shows how the overlap between UniProtKB and the sets derived from the GNF atlas alone (panel A) and the combined transcriptomics and proteomics data (panel B), respectively.

*http://dx.doi.org/10.6084/m9.figshare.1405673*

**Figure S3. Score calibration.** The figure shows that after score calibration, the same confidence score corresponds to the same quality irrespective of the source of the evidence.

*http://dx.doi.org/10.6084/m9.figshare.1405674*

